# Impact of Butyrate on Small and Large Airways: Effects on Cell Viability, Inflammatory Changes and Permeability

**DOI:** 10.1101/2024.02.05.578991

**Authors:** Abdullah Burak Yildiz, Gizem Tuse Aksoy, Nur Konyalilar, Ozgecan Kayalar, Seval Kubra Korkunc, Hasan Bayram

**Affiliations:** Koc University School of Medicine, Istanbul, Turkey; Koc University Research Centre for Translational Medicine (KUTTAM), Koç University, Istanbul, Turkey; Department of Pulmonary Medicine, Koc University School of Medicine, Istanbul, Turkey

**Keywords:** Butyrate, chronic airway disease, hydrogen peroxide, SCFA, inflammation, permeability, cell viability

## Abstract

Chronic airway diseases, such as Chronic Obstructive Pulmonary Disease (COPD) and asthma pose a significant global health burden. The pathophysiology involves chronic inflammation, with oxidative stress playing a crucial role in disease severity. Current treatments, especially for COPD, have limitations, necessitating exploration of alternative therapeutic approaches. In this study, we investigated the potential effects of butyrate, a short-chain fatty acid, on airway epithelial cells. Human bronchial epithelial cells (BEAS-2B) and bronchiolar epithelial carcinoma cells (A549) were cultured and exposed to hydrogen peroxide (H_2_O_2_) to induce oxidative stress. Butyrate was then applied at various concentrations, and the impact on cell viability, epithelial permeability, inflammatory cytokines, and gene expression was assessed. Our cell viability experiments revealed a dose-dependent reduction in viability with H_2_O_2_, while butyrate was found to be safe as it did not affect cell viability. Additionally, butyrate showed decrease in small airway permeability. Butyrate demonstrated anti-inflammatory properties, suppressing H_2_O_2_-induced release of interleukin (IL)-6, IL-8, and granulocyte macrophage colony-stimulating factor (GM-CSF) in large airways. Gene expression analysis further highlighted complex regulatory effects of butyrate on inflammatory pathways. Our study suggests that butyrate may have potential therapeutic benefits in chronic airway diseases by modulating inflammation, permeability, and gene expression. However, further research, including in vivo studies and exploration of endogenous butyrate utilization, is needed to fully understand its pharmacodynamics and clinical relevance. Our findings contribute to the understanding of short-chain fatty acids as potential candidates for respiratory disease treatment.

## INTRODUCTION

Chronic airway diseases, including Chronic Obstructive Pulmonary Disease (COPD) and asthma, are a significant cause of death and morbidity.(Agustí, Celli et al. 2023, Levy, Bacharier et al. 2023) Approximately 600 million people globally suffer from COPD, and it annually causes 3 million deaths worldwide.(Rabe and Watz 2017, Agustí, Celli et al. 2023) Asthma affects 1-18% of the global population and is a major public health concern.(Levy, Bacharier et al. 2023)

COPD is a chronic inflammatory condition that affects the airways and lung parenchyma, and the main risk factors for the disease are tobacco smoke and air pollutants and other inhaled toxicants.(Vogelmeier, Criner et al. 2017) Inflammation in COPD is caused by the recruitment and activation of immune cells such as CD8+ lymphocytes, neutrophils, and macrophages (Hogg et al., 2004), as well as the release of inflammatory mediators.(Vogelmeier, Criner et al. 2017) This leads to a protease-antiprotease imbalance which is associated with increased oxidative stress at cellular level, decreased antioxidant capacity (Malhotra et al., 2008), and protease-antiprotease imbalance.(Vogelmeier, Criner et al. 2017) On the other hand, asthma is characterized by increased airway hyperreactivity, and airway inflammation, which is dominated in large airways.(Levy, Bacharier et al. 2023) Structural cells, like airway epithelial cells(Bayram, Devalia et al. 1998), and immune cells such as eosinophils and lymphocytes, contribute to the disease development, as do inflammatory mediators released from these cells. (Levy, Bacharier et al. 2023)

Reactive oxygen species (ROS) and reactive nitrogen species (RNS) also contribute to inflammation and disease severity in asthma.(Raby, Michaeloudes et al. 2023) The oxidative and nitrosative stress activate pathways like mitogen- activated protein kinase (MAPK) and nuclear factor kappa-light-chain-enhancer of activated B cells (NF-κB) and increases the expression of inflammatory mediators and cell death regulating proteins that could lead to inflammatory changes, as well as cell apoptosis. (Barnes 2008, Gogebakan, Bayraktar et al. 2014) Although the inflammation in asthma is sensitive to corticosteroids, the effects of these drugs in COPD is limited. While bronchodilators may provide symptom relief (Barnes, Ito et al. 2004) in COPD, steroids do not effectively treat the underlying inflammation, and there are currently limitations in the treatment and the disease progression in COPD. (Barnes 2008, Vogelmeier, Criner et al. 2017)

The airway epithelium helps protect the airways by serving as a barrier that prevents the passage of harmful substances in inhaled air to the deeper layers of the airway wall(Bayram, Devalia et al. 1998, Aghapour, Raee et al. 2018, Akdis 2021). This integrity is maintained by intercellular junction complexes such as tight junctions and desmosomes that connect neighboring cells.(Nishida, Brune et al. 2017) Airway epithelial cells also orchestrate the airway inflammation by secreting growth factors, chemokines, and cytokines as a requisite of their metabolic functions.(Knight and Holgate 2003)

Studies have shown that oxidative stress, increased airway epithelial permeability, and inflammatory changes with apoptosis are common features of asthma and COPD(Kirkham and Rahman 2006, Nishida, Brune et al. 2017). Indeed, exposure to oxidant gases like ozone and nitrogen dioxide (NO_2_) can increase the permeability of bronchial epithelial cells from individuals with asthma and lead to increase in the production of inflammatory mediators like IL-8 and GM-CSF(Bayram, Devalia et al. 1998). In COPD, increased oxidative stress has been linked to inflammation caused by immune cells like macrophages, neutrophils, and T cells, and to the production of inflammatory mediators like TNF-α, IL-1, IL-6, and proteases in epithelial cells with compromised permeability.(King 2015)

Short-chain fatty acids (SCFAs) like acetate, propionate, and butyrate are produced by the fermentation of indigestible carbohydrates by anaerobic bacteria in the intestine.(Yan and Ajuwon 2017) These SCFAs act as regulatory molecules that influence the proliferation, differentiation, and apoptosis of epithelial cells through histone deacetylation and G protein-coupled receptors.(Roduit, Frei et al. 2019) SCFAs can also suppress the Th2 response of the immune system by affecting T cells and dendritic cells(Cait, Hughes et al. 2018).

Butyrate has been shown to increase the activity of adenosine monophosphate-activated protein kinase and accelerate the formation of tight junctions in intestinal epithelial cells, protecting the epithelial barrier. However, high doses of butyrate can be toxic to cells (Peng, Li et al. 2009). In vitro studies suggest that SCFAs may have anti-inflammatory properties and may improve the epithelial barrier integrity by inhibiting the release of mediators that damage the barrier and suppressing the expression of tight junction proteins in intestinal epithelial cells (Chen, Kim et al. 2017). Nevertheless, it is not yet clear if SCFAs can have similar effects on pathological changes in airway epithelial cells.

This study aims to investigate the effect of butyrate on cell viability, epithelial permeability, and inflammatory changes induced by hydrogen peroxide (H_2_O_2_) in airway epithelial cells. The findings of this study may contribute to our understanding of the potential properties of SCFAs in the treatment of chronic airway diseases such as asthma and COPD.

## MATERIAL and METHOD

### Airway Epithelial Cell Culture

Human bronchial epithelial cells (BEAS-2B) and adenocarcinoma human alveolar basal epithelial cells (A549) were cultured in RPMI 1640 (Gibco, Thermo Fisher Scientific, USA) and DMEM medium containing 10% fetal bovine serum (FBS) and 1% penicillin/streptomycin (Corning, Thermo Fisher Scientific, USA), respectively. Cells were grown on 75 cm^2^ cell culture flask at 37 °C in a 5% CO_2_ humidified incubator.

### Incubation of Airway Epithelial Cell Cultures with Hydrogen peroxide (H_2_O2) and Butyrate

The cells were seeded on 6-well (3×10^5^ cells/well) and 24-well (5×10^4^ cells/well) culture plates for qPCR and cell viability experiments, respectively. BEAS-2B and A549 cells were plated in 24 well plates. Two days later, the growth medium was replaced with serum-deprived (SF) medium consisting of RPMI-1640, 0.1% FBS, and 1% penicillin/streptomycin, and incubated for one day. Moreover, confluent cells were exposed to various concentrations of H_2_O_2_ (0, 50, 100 and 200µM) for 24 hours or different concentrations of sodium butyrate (0, 0.3, 1, and 3 mM) for 30 minutes. Cell culture supernatant was collected and stored at - 80°C, and analysis of IL-8, IL-6, and GM-CSF performed by ELISA. Additional set of cultures were established in 24-well trans well inserts for transepithelial-endothelial resistance (TEER) experiments.

### Measurement of Cell Viability with 3-(4,5-dimethylthiazol-2-yl)-2,5-diphenyltetrazolium bromide (MTT)

At the end of treatment protocol, cell viability was measured by 3-(4-5- dimethylthiazol-2-yl)-2,5- diphenyltetrazolium bromide (MTT, 1mg/ml) as described previously (Bayram, Fakili et al. 2013). The absorbances of the cells were quantified in 570 nm using a microplate reader (Biotek USA). Following medium aspiration, the cells were then washed with phosphate-buffered solution (PBS). MTT incubation and absorbance, and the determination of cell viability percentage were performed as previously described. cell viability was measured by 3-(4-5-dimethylthiazol-2-yl)-2,5- diphenyltetrazolium bromide (MTT, 1mg/ml) as described previously. (Bayram, Fakili et al. 2013)The absorbances of the cells were quantified in 570 nm by using a microplate reader (Biotek USA).

### Analysis of IL-6, IL-8, and GM-CSF

The determination of interleukin (IL)-6, IL-8, and granulocyte macrophage -colony stimulating factor (GM-CSF) were analysed by enzyme-linked immunosorbent assay (ELISA, R&D System Duoset, USA) according to the manufacturer’s instructions.

### Analysis of Relative mRNA Expressions of ERK5, NF-κB, c-Jun, IL-6, IL-8, GM-CSF, and p21 with RT-qPCR

Total RNA was isolated from BEAS-2B and A549 cells (2×10^5^ cells/ well) seeded to 6-well plate by using an RNA isolation kit according to the manufacturer’s instructions (Zymo, USA). Complementary DNA (cDNA) was synthesized via cDNA synthesis kit (Biorad, USA) and target genes included ERK5, NF-κB, c-Jun, IL-6, IL-8, GM-CSF, and p21 were amplified with Light Cycler Syber Green Master Mix (Roche, Germany) on Light Cycler 480 instrument. Glyceraldehyde-3-phosphate dehydrogenase (GAPDH) was used as a housekeeping gene. Relative gene expression compared to control was presented as 2^-ΔΔCt^= (ΔΔC_t_= ΔC_t treated_ - ΔC_t control_)(Kayalar, Oztay et al. 2020). Primers are listed in Table 1.

**Table 1.**
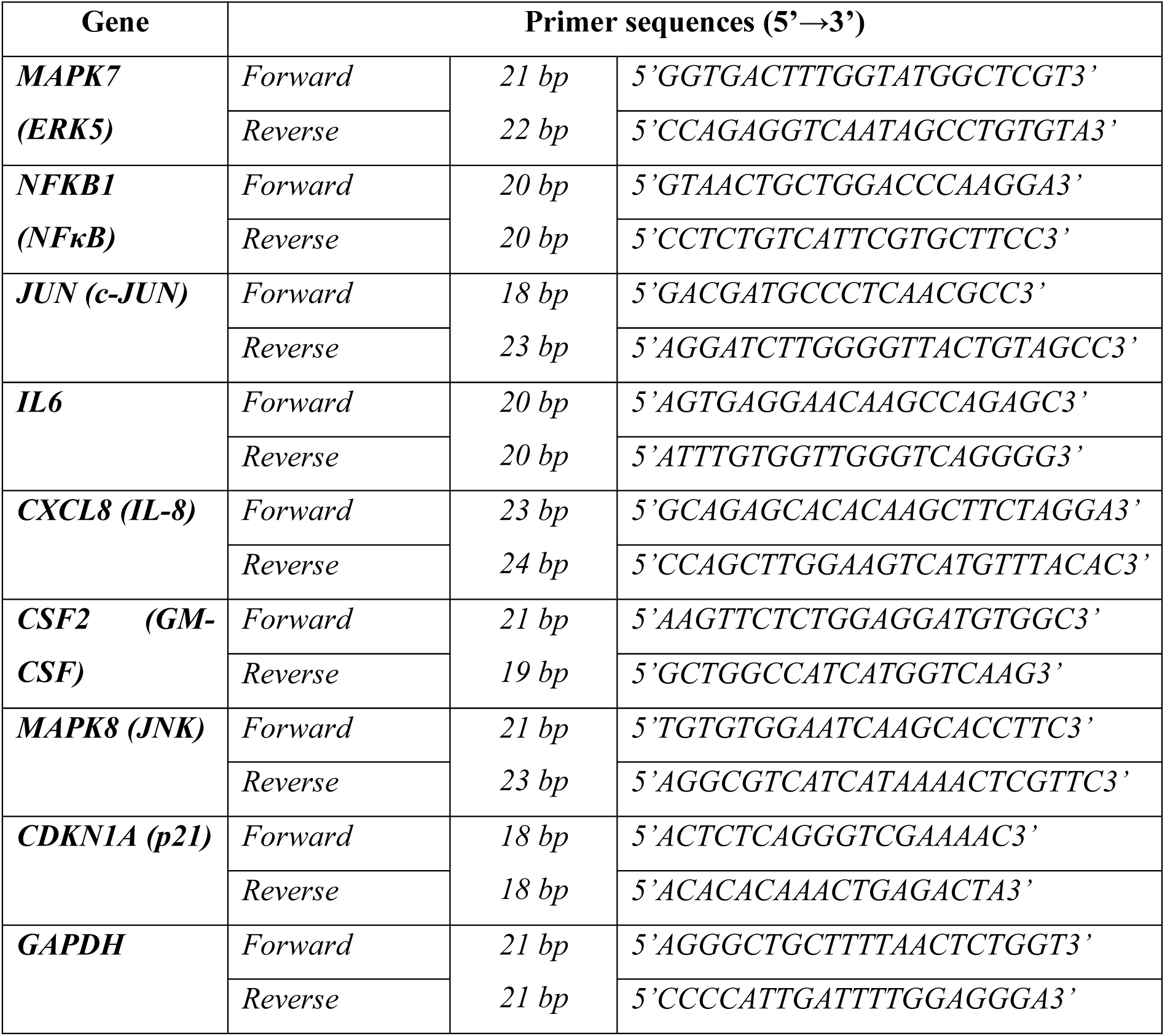
Primers used for real-time qPCR.

### Measurement of Transepithelial electrical resistance (TEER) in BEAS-2B and A549 Cells Incubated with Butyrate and H_2_O_2_

BEAS-2B and A549 cells were plated in 24 well inserts (3×10^4^ cells/insert). After the treatment protocol, TEER was measured by using EVOM5 TEER device at 2^nd^ hours, 4^th^ hours, 6^th^ hours, and 24^th^ hours.

### Statistical Analysis

The normality of the data was assessed using the D’Agostino and Pearson omnibus normality test, continuous variables were conducted employing one-way variance analysis (ANOVA) / Dunnett’s multiple comparison tests or Kruskal–Wallis with Dunn’s multiple comparison tests. Results are presented in terms of means ± SEM or medians ± interquartile ranges (Q1 and Q3) alongside lower and upper ranges. p ≤ 0.05 was regarded as significant. Statistical analyses were executed using PRISM version 8 (GraphPad Software Inc., San Diego, CA, United States).

## RESULTS

### Cellular Viability

#### Butyrate Did Not Prevent BEAS 2B Cell Viability Decreased by H_2_O_2_

The investigation aimed to assess the cytotoxic impact of H_2_O_2_ and butyrate on BEAS-2B cells through varying concentrations. H_2_O_2_ concentrations of 10 µM, 25 µM, 50 µM, 100 µM, and 200 µM were utilized, revealing a dose-dependent reduction in cell viability. Notably, 10 µM and 25 µM concentrations exhibited limited impact on cell viability, while 50 µM, 100 µM, and 200 µM resulted in a notable decline **(Fig. 1a).** Consequently, the optimal concentration was determined as 100 µM. Subsequently, cells were exposed to varied concentrations of butyrate (0.3 mM, 1 mM, and 3 mM) in combination with the identified optimal H_2_O_2_ concentration. None of the doses of butyrate was found toxic to cells and co-treatment did not significantly affect cell viability **(Fig. 1b).**

**Figure 1:**
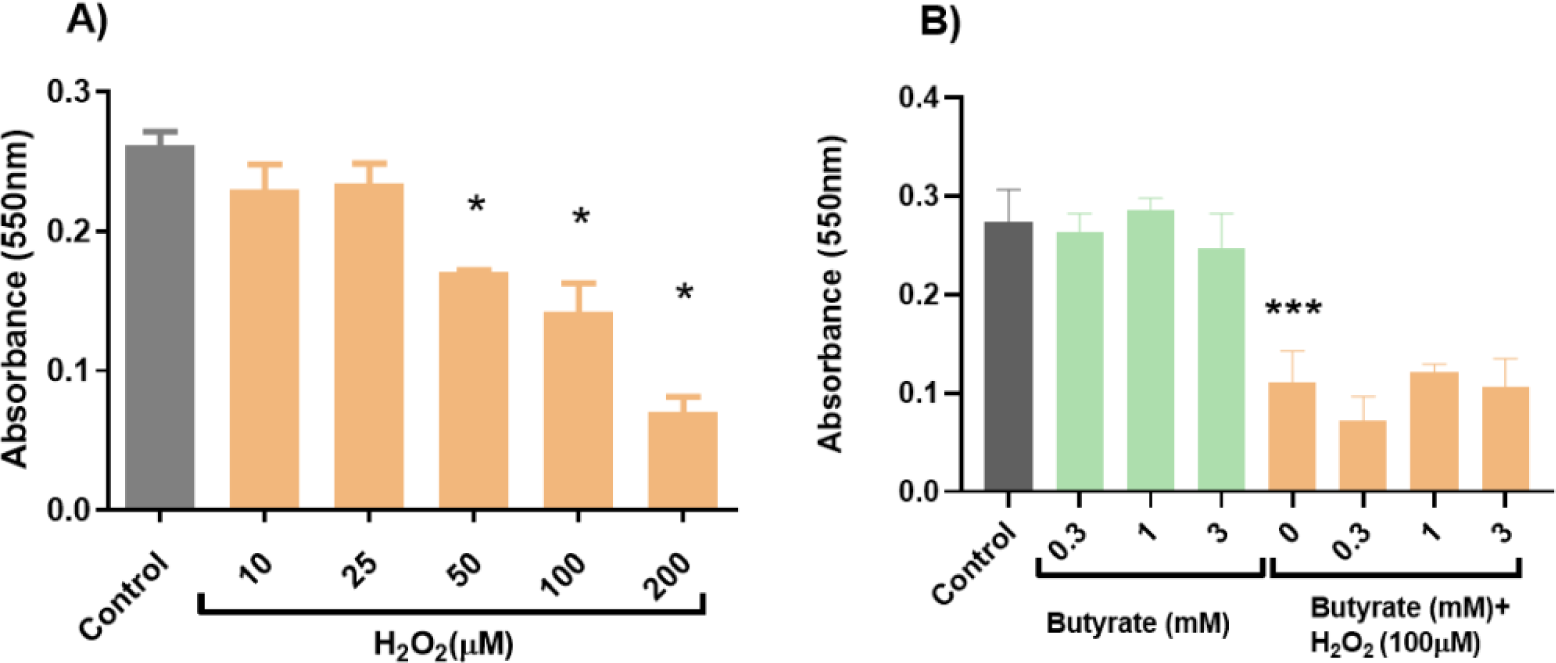
**(a)** Effect of H_2_O_2_ (10-200 μM) on BEAS-2B cell viability at 24 h. **(b)** Effect of 0.3, 1, and 3mM butyrate pre-treatment in the absence and presence of H_2_O_2_ on BEAS-2B cell viability at 24 h *p<0.05, ***p<0.001 compared to control group

#### Butyrate Did Not Prevent A549 Cell Viability Decreased by H_2_O_2_

The investigation aimed to assess the cytotoxic impact of H_2_O_2_ and butyrate on A549 cells, reflecting butyrate’s effect on small airways physiology. H_2_O_2_ concentrations of 100 µM, 300 µM, and 600 µM were utilized, revealing a reduction in cell viability at all doses. Notably, 100 µM, 300 µM, and 600 µM resulted in a prominent decline in cell viability in both 24h and 48h measurements **(Figure 2a),** whereas lower doses were found to be less effective. The optimal concentration was determined as 600 µM for use in further experiments.

**Figure 2:**
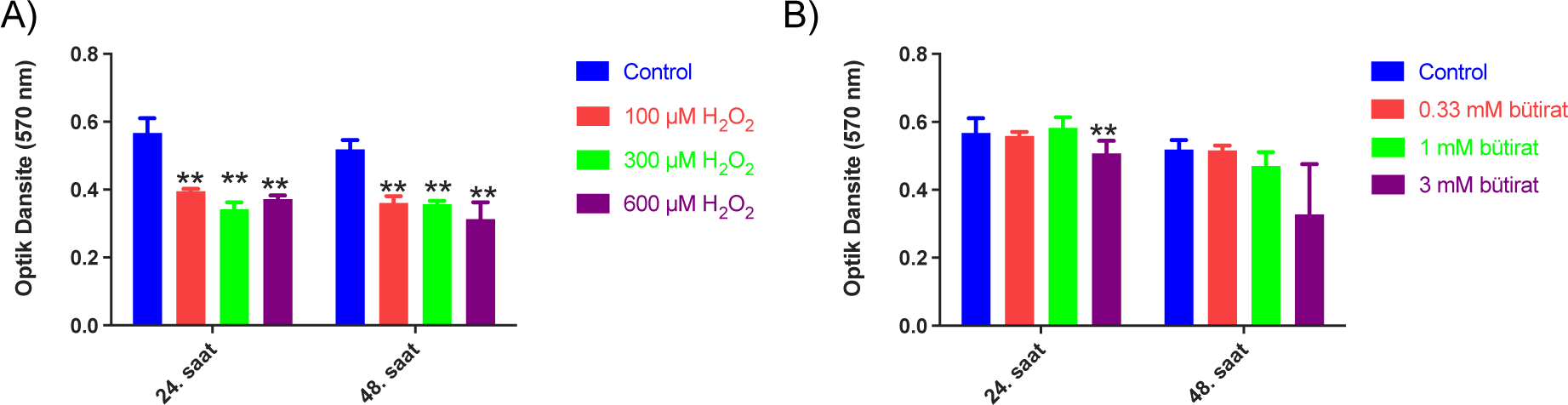
**(a)** Effect of H_2_O_2_ (100-600 μM) on A549 cell viability at 24 and 48 h. **(b)** Effect of 0.33, 1, and 3mM butyrate on A549 cell viability at 24 and 48h. **\*\***p<0.01 compared to control group.

Butyrate showed toxicity at a 3 mM dose in 24h **(Figure 2b),** prompting subsequent experiments to use cells exposed to butyrate concentrations of 0.11 mM, 0.3 mM, and 1 mM, in combination with the 600 µM H_2_O_2_ concentration. Co-treatment did not significantly affect cell viability **(Figure 3)**.

**Figure 3:**
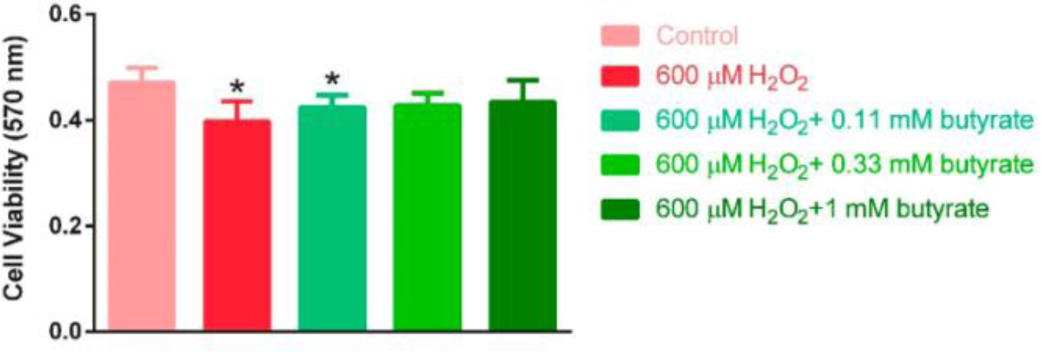
Effect of 0.11, 0.33, and 1 mM butyrate pre-treatment in the presence of 600 μM H_2_O_2_ on A549 cell viability at 24 h. **\***p<0.05 compared to control group.

### Cellular Permeability

#### Butyrate Had No Effect on H_2_O_2_ increased BEAS 2B Cellular Permeability

At 24 hours, 100 µM H_2_O_2_ demonstrated a decrease in TEER at 24h, whereas 1mM butyrate decrease TEER at 2h compared to the control group in BEAS 2B. **(Figure 4).** Butyrate has no effect with co-treatment on TEER measurements after H_2_O_2_ decreased TEER completely.

**Figure 4:**
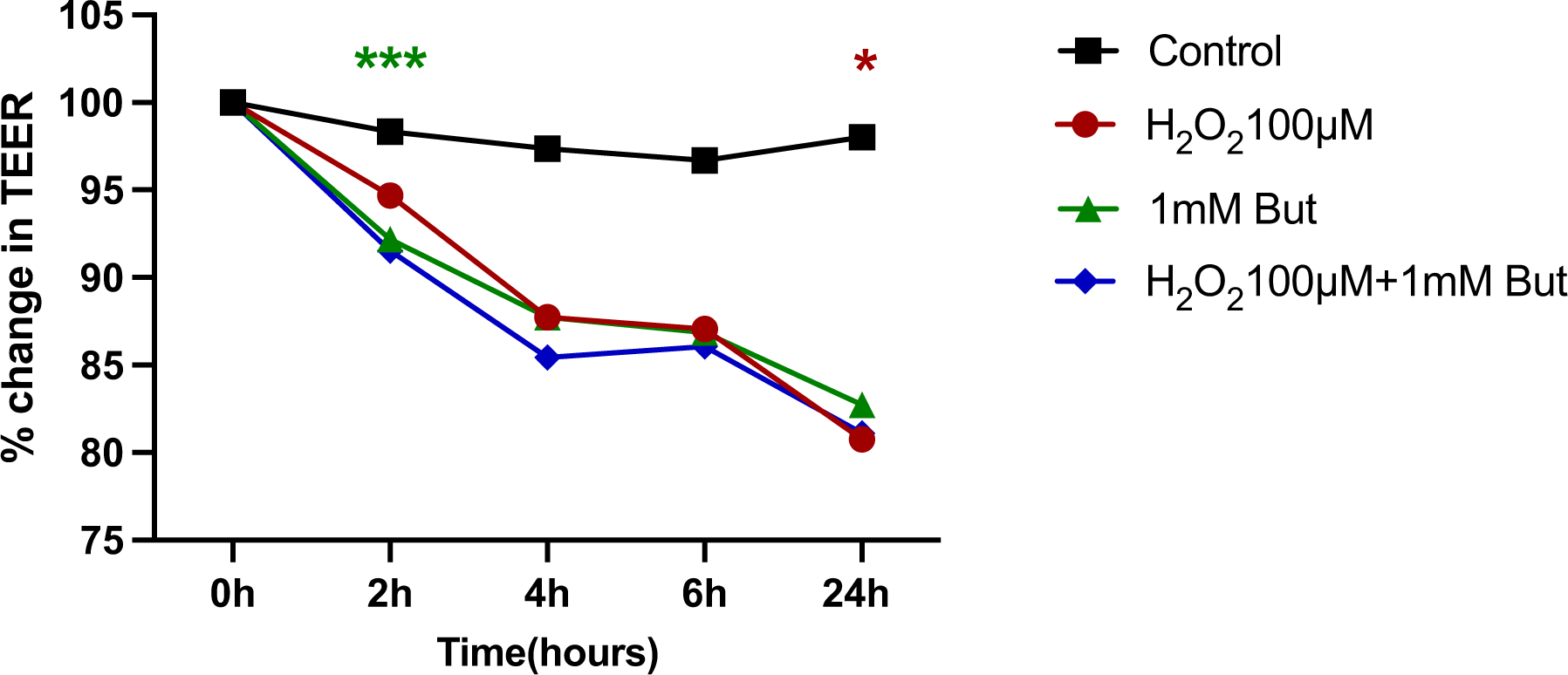
TEER at 0, 2, 4, 6, and 24h of butyrate and/or H_2_O_2_ treatment in BEAS 2B cells; H_2_O_2_ **\***p<0.05 compared to control group; \*\***\*** Butyrate p<0.001 compared to control group

### Inflammatory Cytokines

#### Butyrate Supresses H_2_O_2_ Induced IL-6, IL-8 and GM-CSF Levels in BEAS 2B Cells

The impact of varying concentrations of H_2_O_2_ (10 µM, 25 µM, 50 µM, 100 µM, and 200 µM) on inflammatory cytokines (IL-6, IL-8, and GM-CSF) in BEAS-2B cells was assessed with ELISA. Subsequent experiments involved treating BEAS-2B cells with different concentrations of butyrate for 30 minutes, followed by exposure to 100 µM H_2_O_2_ **(Figure 5).**

**Figure 5:**
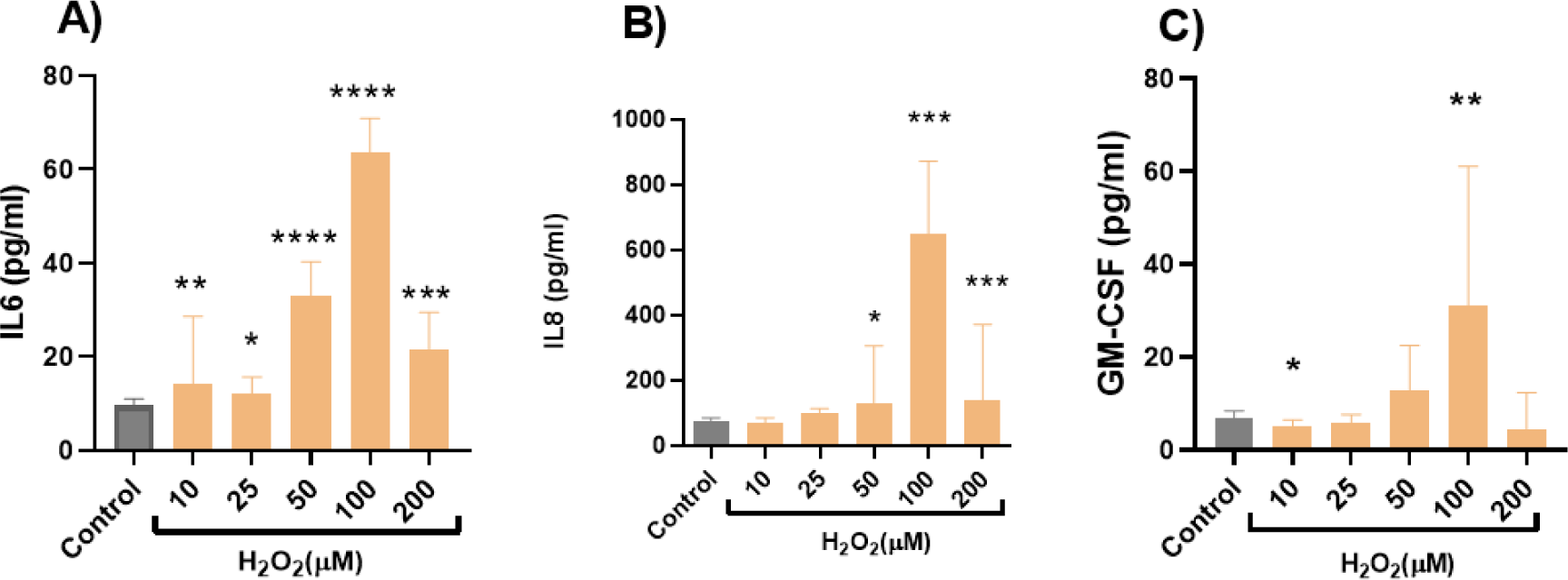
**(a)** Effects of 10, 25, 50, 100, and 200 μM H O treatment on levels of IL-6; **(b)** Effects of 10, 25, 50, 100, and 200 μM H O treatment on levels of IL-8; **(c)** Effects of 10, 25, 50, 100, and 200 μM H O treatment on levels of GM-CSF; *p<0.05, **p<0.01, ****p<0.0001 compared to the control

The analysis revealed significant increases in all concentrations of IL-6 compared to the control group, with 100 µM demonstrating the highest elevation (p<0.0001) **(Figure 5a).** Notably, 0.33 mM butyrate exhibited a significant decrease in IL-6 release, while other concentrations showed no effect. When combined with H_2_O_2_ (100 µM), 0.33 mM butyrate further reduced IL-6 levels. However, 1 mM and 3 mM butyrate with H_2_O_2_ treatment elevated IL-6 release compared to the H_2_O_2_ treatment alone group **(Figure 6a).**

**Figure 6:**
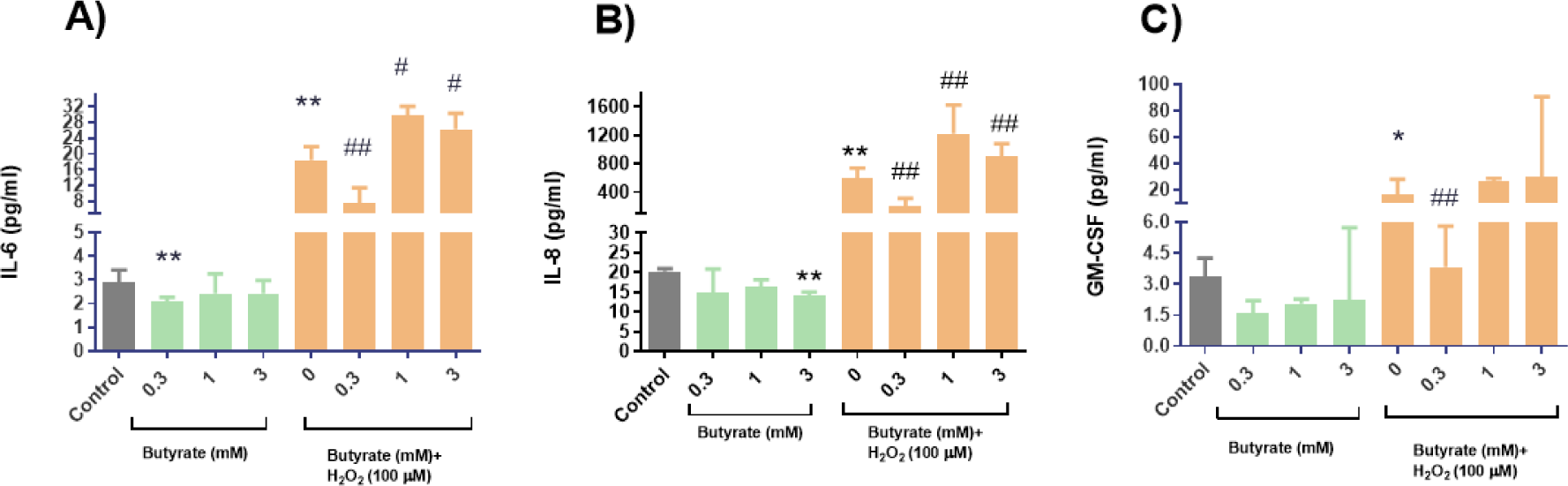
Changes of inflammatory cytokines of 0.3, 1, and 3mM butyrate pre-treatment in the absence and presence of H_2_O_2_ on BEAS-2B cell. **(A)** IL-6, **(B)** IL-8, **(C)** GM-CSF *p<0.05, **p<0.01 compared to the control; #p<0.05, ##p<0.01 compared to 100 uM H_2_O_2_

IL-8 release remained unaffected by some H_2_O_2_ concentrations (10 µM and 25 µM). However, 100 µM H_2_O_2_ notably increased IL-8 release compared to 50 µM and 200 µM **(Figure 5b).** Butyrate treatment independently decreased IL-8 release significantly at 3 mM compared to the control group. Results of butyrate treatment with H_2_O_2_ mirrored those observed for IL-6 **(Figure 6b).**

Furthermore, 10 µM H_2_O_2_ exhibited a significant decrease in GM-CSF release, while 100 µM H_2_O_2_ led to increased GM-CSF release akin to IL-6 and IL-8 experiments. Independent butyrate treatment decreased GM-CSF release, with the combined treatment with H_2_O_2_ demonstrating similar patterns to IL-6 and IL-8 experiments compared to the H_2_O_2_ group **(Figure 6c).**

### mRNA expression

#### mRNA Expression Results in BEAS 2B Cells

Our investigation into the gene expression dynamics of NF-κB, c-JUN, ERK5, p21, JNK1, IL-6, IL-8, and GM-CSF, utilizing RT-qPCR analysis, elucidated distinct responses to both butyrate and its combination with H_2_O_2_ treatment (100 µM).

NF-κB exhibited a notable elevation in expression levels following exposure to H_2_O_2_ and 0.3 mM butyrate, demonstrating significant increases compared to the control group. However, 1 mM and 3 mM butyrate co-administered with H_2_O_2_ displayed no discernible effects on NF-KB expression, with 0.3 mM showing a notable rise compared to the H_2_O_2_ group. (**Figure 7a**) c-JUN expression was prominently increased in the H_2_O_2_ group compared to the control, while significant alterations were observed across all concentrations of butyrate combined with H_2_O_2_ (100 µM), particularly notable with 0.3 mM butyrate in conjunction with H_2_O_2_. **(Figure 7b)** ERK5 expression remained unaltered following the different treatment regimens. **(Figure 7c)**

**Figure 7:**
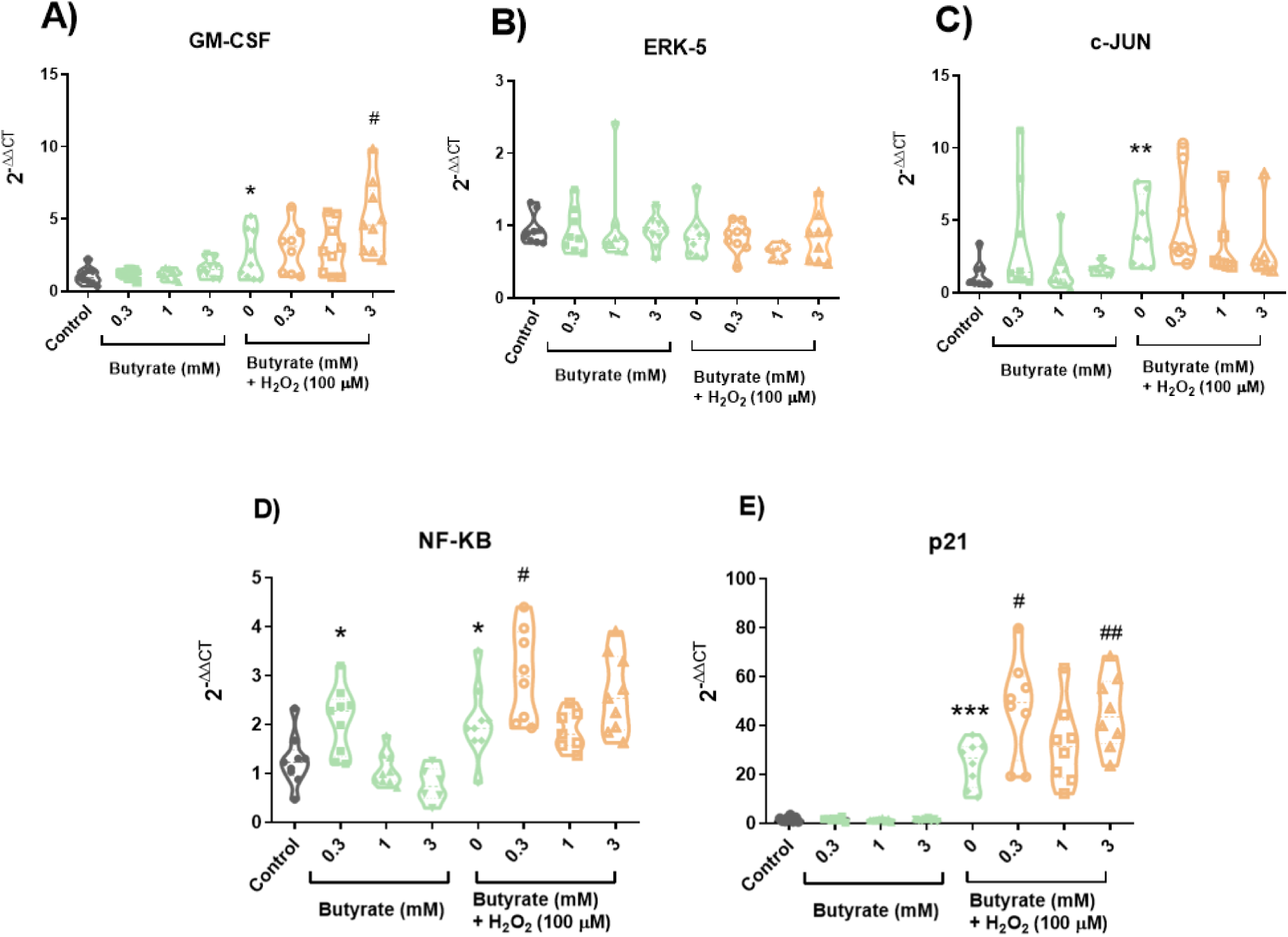
Changes of gene expressions of GM-CSF **(A),** ERK5 **(B),** c-JUN **(C),** NF-κB **(D),** and p21 **(E)** respectively in BEAS 2B cells. *p<0.05, **p<0.01, ***p<0.001 compared to the control; p<0.05, ##p<0.01 compared to 0

The influence on p21 expression was apparent in co-treatment, whereas the impact was less evident with sole 0.3 mM and 3 mM butyrate treatments. However, heightened p21 expressions were noted with H_2_O_2_ treatment, further augmented when combined with butyrate. **(Figure 7d)** IL-6 expression notably increased with 0.3 mM and 3 mM butyrate treatments, whereas 1 mM butyrate alone did not elicit significant alterations. Additionally, the H_2_O_2_ group demonstrated a marked increase in IL-6 expression. Interestingly, while significant rises were observed at 0.3 mM and 3 mM in conjunction with H_2_O_2_ (100 µM), no substantial difference was noted at 1 mM butyrate combined with H_2_O_2_ compared to the H_2_O_2_ group. **(Figure 7e)** IL-8 expression remained relatively unaffected by sole butyrate treatments yet displayed notable increases following exposure to H_2_O_2_. Furthermore, IL-8 expressions were heightened across all concentrations of butyrate compared to the control group. **(Figure 7f)** GM-CSF expressions were increased in response to H_2_O_2_, with a more pronounced increase observed in the presence of 3 mM butyrate combined with 100 µM H_2_O_2_. **(Figure 7g)**

These findings highlight varied regulatory effects of butyrate and its combination with H_2_O_2_ on gene expression, underscoring their complex interplay in cellular responses.

#### mRNA Expression Results in A549 Cells

Our investigation into the gene expression dynamics of NF-κB, c-JUN, ERK5, p21, IL-8, utilizing RT-qPCR analysis, elucidated distinct responses to both butyrate and its combination with 600 µM H_2_O_2_ treatment.

Regarding IL-8, 600 µM H_2_O_2_ treatment showed a significant increase in IL-8 mRNA expression, which was attenuated by 0.33- and 1-mM concentrations of butyrate **(Figure 8a).** c-Jun mRNA expression was increased with 600 µM H_2_O_2_ compared to the control, whereas pre-treatment with butyrate significantly decreased this augmented c-Jun mRNA expression in all doses, exhibiting the highest significant change in the 1 mM butyrate treatment **(Figure 8b)**. Examination of ERK-5 revealed that pre-treatment with 0.33 mM butyrate led to a decrease in ERK-5 mRNA expression, while no significant results were detected with other doses **(Figure 8c)**. No significant changes were observed in NF-κB and p21 mRNA expression in small airway epithelial cells in vitro **(Figure 8d, e)**.

**Figure 8:**
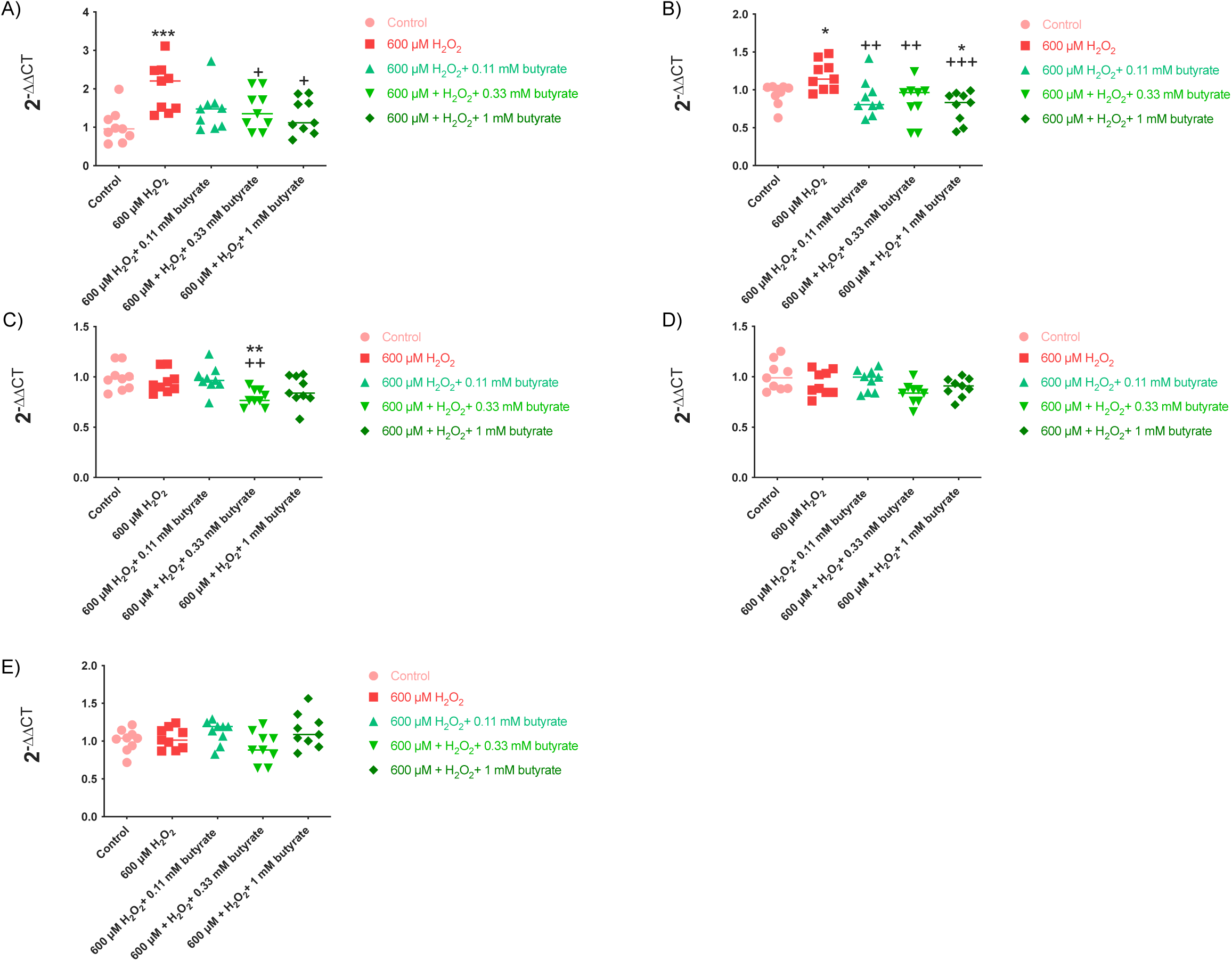
Effect of 0.11, 0.33 and 1 mM butyrate pre-treatment on gene expressions of IL-8 **(A)**, c-Jun **(B)**, ERK5 **(C),** NF-κB **(D)**, and p21 **(E)** with 600 μM H_2_O_2_ in A549 cells in qPCR. **\***p<0.05, \***\***p<0.01, \*\***\***p<0.001 compared to control group. +p<0.05, **++**p<0.01, +++p<0.001 compared to 600 μM H_2_O_2_.

## DISCUSSION

This study aimed to explore how butyrate influenced the effects of H_2_O_2_ on cell viability, epithelial permeability, and inflammatory responses in large and small airways. The investigation revealed that butyrate pre-treatment did not prevent H_2_O_2_-induced cytotoxicity in large and small airway cells in vitro, but it demonstrated differential effects on permeability, inflammatory cytokines, and gene expression in these distinct airway cell types, emphasizing the complex and context-dependent regulatory role of butyrate.

By the means of studying butyrate’s effect on small airways with the established cell line A549- adenocarcinomic human alveolar basal epithelial cells(Smith 1977), and with the large airways BEAS 2B nontumorigenic immortalized human bronchial epithelium (Albright, Jones et al. 1990), helped to predict the extent of butyrate and its possible effects in different compartments of lung physiology.

H_2_O_2_, a well-established agent that causes oxidative stress and release of reactive oxygen species has been shown to capture the similar phenotypes regarding cell viability, permeability and inflammatory markers in our lung epithelial cell lines (all figures, (Averill-Bates 2023)). H_2_O_2_ is expected to augment signaling of MAPK and phosphoinositide 3-kinase (PI3K)/protein kinase B (Akt), hypoxia-inducible factor 1α (HIF-1α), NF-κB and other inflammatory pathways similarly established in our experiments in protein and mRNA expression (Figure 4-7, (Averill-Bates 2023)) and can lead to apoptosis of cells **(Figure 1a and 2a)**

Within the recent decade, SCFAs such as acetate, propionate, and butyrate are found to be fundamental compounds present in the body, serving as physiological regulatory molecules involved in the proliferation, differentiation, and apoptosis of epithelial cells (Loius et al., 2014). Literature includes studies on gastrointestinal epithelial cells, Caco-2 cells, showing a dose dependent butyrate effect similarly observed in our study, with its high doses having cellular toxicity in A549 cell line ((Peng, Li et al. 2009), **Figure 2b**), however same doses of butyrate have shown no effect in cellular toxicity in Beas 2B cell lines **(Figure 1b),** indicating butyrate could be safe to use as a supplement for respiratory diseases.

We speculate that toxicity with high dose seen in A549 cell line is due to cell specificity. Since butyrate has a potential anti-tumor effect (Celasco, Moro et al. 2014, Dos Santos, de Farias et al. 2014), using A549 lung adenocarcinoma cell line alone, could have misrepresented the actual effect of cell viability at higher doses, leading to its anti-tumor effects to be more pronounced similar to Caco-2 cell line which is derived from colorectal adenocarcinoma (Peng, Li et al. 2009). Even though BEAS 2B cell line is immortalized by adenovirus transfection, tumorigenic properties are less pronounced. (Albright, Jones et al. 1990). This needs to be confirmed by in vivo studies to supplement butyrate in mice for respiratory toxicity.

H_2_O_2_ demonstrated a decrease in TEER reflecting a cellular permeability increase which was not prevented by butyrate. This contrasts with the limited in vitro studies conducted on butyrate and other SCFAs, indicating that SCFAs predominantly suppress the formation of inflammation in intestinal epithelial cells, thereby inhibiting the release of mediators that lead to barrier breakdown and rescue of the suppressed tight junctions (Chen, Kim et al. 2017). Interestingly, butyrate alone has increased cellular permeability at 2h. We believe careful consideration on cellular permeability should be done as TEER is not a sensitive method, further work is needed to check these effects in a simultaneous use of universally acknowledged permeability measurement method, dextrane passage assay (Elamin, Masclee et al. 2013) with TEER, should be employed in A549 and BEAS 2B cells to strengthen the results. Further, tight junctions and barrier components can be checked in detail.

Butyrate is found to supress H_2_O_2_ mediated IL-6, IL-8 and GM-CSF levels in large airways in vitro **(Figure 5 and 6)** correlated with its potential anti-inflammatory effects in the literature (Chen, Kim et al. 2017). This study is the first study to suggest butyrate as a potential antioxidant and anti-inflammatory agent, shows promising results in respiratory airway cells and these effects are visible in cytokine level.

Going back one more step, we wanted to see if there are any changes in transcriptomic level with butyrate. H_2_O_2_ has shown upregulation or nonspecific trend in upregulation in transcription of inflammatory pathways **(Figure 7)**, butyrate did not suppress this effect, interestingly, butyrate has shown to have increase some inflammatory genes (butyrate alone: NF-κB; co-treatment: NF-κB, p21, GM-CSF). In small airways, H_2_O_2_ have upregulated some (IL-8 and c-Jun) but not all inflammatory genes. IL-8, c-Jun and ERK5 were suppressed with butyrate co-treatment in the presence of H_2_O_2_, whereas NF-κB and p21 show no difference. **(Figure 8)**

These findings emphasize the regulatory effects of butyrate before H_2_O_2_ exposure on gene expression in most of the inflammatory pathways, highlighting butyrate as a potential anti-inflammatory agent effective in small airways, including its associated changes in transcriptomic level.

There is a need to address why some genes do not exhibit parallel results in both small and large airways, possibly due to heterogeneity between replicated experiments. There could be epigenetic changes which we could check with ATACseq, transcriptomic chances of other inflammatory genes with RNAseq, or on human small and large airways cultured in vitro and conducting further functional studies in vitro. However, these aspects are beyond the scope of this manuscript.

Even though this study is the first to mention the possible beneficence of butyrate for asthmatics and COPD patients by it being safe and non-toxic, inflammatory cytokines, more studies in respiratory system needs to be conveyed and more complex systems are needed to represent a closer COPD and asthma model. We believe SCFAs are important mediators that could be used for many systems and its known antioxidant nature makes it a potential candidate to target and treat COPD and asthma patients as both conditions have oxidative stress and its effects as primary drivers of pathogenesis.

This study lacks the insight of endogenous utilization and effects of butyrate and lacks the association of non-epithelial cells and microenvironment with SCFAs. Contrary to the lack of in vivo studies, checking the desired effects in two common cell lines for both large and small airways, BEAS 2B and A549, strengthened our report. Oxidative stress, considered a primary driver in the pathogenesis of both asthmatic/COPD patients and in the cellular effects of H_2_O_2_ mimicking disease progression, serves as the central point of investigation. To get a broader view of butyrate’s effect on large and small airways and correlate anti-inflammatory and antioxidant effects, conducting a bulk RNA sequencing for our butyrate-treated BEAS 2B and A549 cell lines would have bolstered our hypothesis.

Another problem to think about is the actual pharmacodynamics of SCFA when taken by a living organism. Most of the studies concentrate on gastrointestinal (GI) system due to its highly known regulation and continuous utilization by GI microbes and environment. Even though the half-life of butyrate when given intravenously to acute leukemic patients is known to have an half-life of 6-7 minutes (Miller, Kurschel et al. 1987), the actual distribution within lungs when given exogenously is question to be answered before claiming its possible beneficial effects in asthma and COPD patients.

To sum up, even though this paper does not include mechanistic insight and possible interactions with non-epithelial cells and microenvironment in the lung, it shows promising results on butyrate for its use in exogenous utilization and a as possible agent that could be a beneficial substance to provide for patients with COPD or asthma. Results indicate that butyrate has anti-inflammatory effects, making it a promising substance that could be used in the future. After establishing butyrate’s effect in vivo, further steps would be required to compare and use it as an additive option in experiments along with the gold standard therapies in COPD and asthma.

## Acknowledgements

The authors gratefully acknowledge use of the services and facilities of the Koç University Research Center for Translational Medicine (KUTTAM), funded by the Republic of Türkiye Ministry of Development. Abdullah Burak Yildiz was supported by TUBITAK Scholarship.

## Author contributions

Conceptualization: A.B.Y, H.B. Formal analysis: A.B.Y., G.T.A. Investigation: A.B.Y., G.T.A., N.K., O.K., S.K.K. Writing—original draft: A.B.Y. Writing—review & editing: A.B.Y., H.B. Supervision: H.B.

